# Whole Genome Sequencing of 5 Tibetan Sheep Breeds Identifies Selective Signatures to Adaptability at Different High-Altitude Areas in Qinghai-Tibetan Plateau

**DOI:** 10.1101/2020.06.09.141903

**Authors:** Lei-Lei Li, Shi-Ke Ma, Wei Peng, You-Gui Fang, Hong-Yun Fu, Gong-Xue Jia

**Author notes:** These authors contributed equally to this work (Lei-Lei Li, Shi-Ke Ma).

## Abstract

Tibetan sheep is one of primitive Chinese sheep breeds, which achieved the divergence about 2500 years ago in Qinghai plateau region. According to different geographic conditions, especially altitudes, Tibetan sheep evolved into different breeds. In this study, we performed pooled whole genome resequencing of 125 individuals from 5 representative Tibetan sheep breeds. Comparative genomic analysis showed that they can be divided into different clades with a close genetic relationship. However, some genes with common selective regions were enriched for hypoxic adaptability in different breeds living at higher altitude, including *GHR, BMP15* and *CPLANE1*. Furthermore, breed-specific selective regions about physical characteristics, especially wool growth, were found in genes such as *BSND, USP24, NCAPG* and *LCORL*. This study could contribute to our understanding about trait formation and offer a reference for breeding of Tibetan sheep.

## INTRODUCTION

Tibetan sheep is one of representative herbivores in northwestern China, with more than 23 million individuals distributed throughout the Qinghai-Tibet plateau (statistics in 2008) (Du 2011). After the long course of evolutionary history, Tibetan sheep has been adapted to the harsh environment and plays an irreplaceable role in economic and social development for local people. Genomic reconstruction results indicate that Tibetan sheep was originated from northern Chinese ancient sheep ∼3100 years ago and achieved the divergence ∼2500 years ago (Zhao et al. 2017; Hu et al. 2019). Since then, a subgroup of Tibetan sheep continued to southwest expand and reached to the central Tibet area ∼1300 years ago, while the rest of them colonized different areas of Qinghai and gradually evolved into different breeds depending on geographic conditions (Hu et al. 2019).

Qinghai has a varied and complicated topography with altitudes ranging from 1650 m to 6860 m, including eastern Hehuang valley region, southeastern Huangnan mountainous region, middle region around Qinghai lake, northern Qilian mountainous region, northwestern Qaidam basin region and southwestern Qingnan plateau region (Zhang 2009). Due to extreme geographic barrier and rare invasion of external populations, most regions reserve abundant and distinctive Tibetan sheep resources. However, because of ongoing regression of ecological environment, the introduction of modern commercial sheep and the lack of effective conservation efforts, indigenous breeds are facing crisis of population decline and genetic characteristic loss. Therefore, it is important to clarify the population structure and genetic diversity of Tibetan sheep to preserve and utilize the genetic materials efficiently.

Hence, we performed pooled whole genome resequencing of 125 sheep from five representative Tibetan sheep breeds in Qinghai. By analyzing single nucleotide polymorphism (SNP) annotation, we aimed to elucidate the genetic relationships among breeds and excavate the candidate genes or variants responsible for adaptability of Tibetan sheep.

## MATERIALS AND METHODS

### Sample collection

Five phenotypically representative Tibetan sheep breeds were chosen in different geographical regions of Qinghai: Valley sheep (referred to as LD) from Hehuang valley region with an elevation of 2000 to 3000 m, Oula sheep (referred to as HN) from Huangnan mountainous region with an elevation of 3500 to 4000 m, Zeku sheep (referred to as ZK) from region around Qinghai lake with an elevation of 3000 to 3500 m, Grassland sheep (referred to as TJ) from Qilian mountainous region with an elevation of 3000 to 4000 m and Zhashijia sheep (referred to as QM) from Qingnan plateau region with an elevation above 4000 m. Whole blood samples were collected from 25 unrelated individuals with similar traits per breed. The detailed sample information is summarized in Figure 1. All experiments in this study were handled in accordance with the requirements of Animal Ethic and Welfare Committee of Northwest Institute of Plateau Biology, Chinese Academy of Sciences.

**Figure 1.**
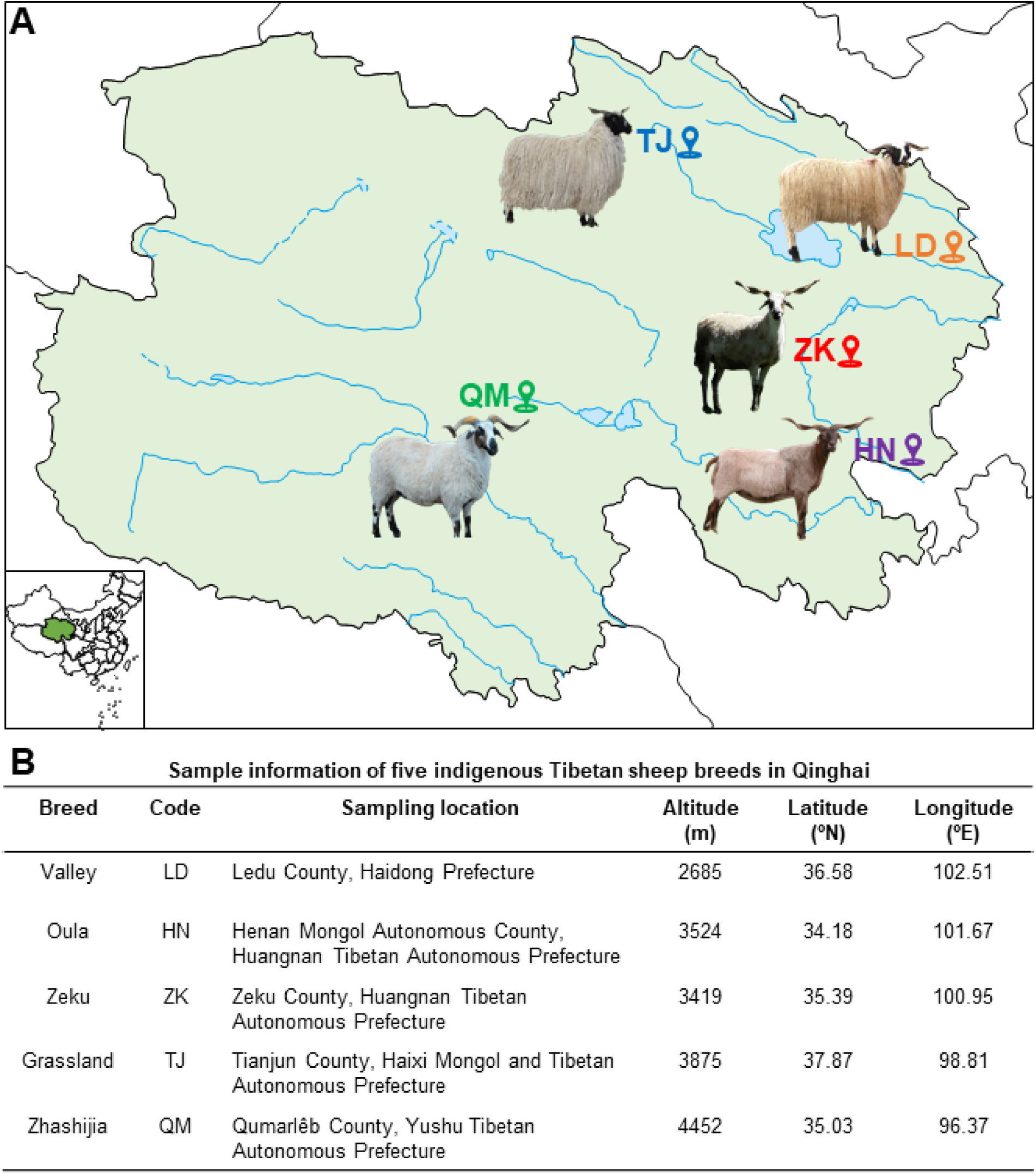
Detailed information of five Tibetan sheep breeds in Qinghai. (**A**) Geographic distribution of five Tibetan sheep breeds in Qinghai. The map was generated using Adobe Illustrator. (**B**) Sample information of five Tibetan sheep breeds in Qinghai, including breed, code, sampling location, altitude (m), latitude (°N) and longitude (°E).

### DNA library construction and resequencing

Genomic DNA was extracted from whole blood samples using a standard phenol-chloroform method (Russell and Sambrook 2001). After dilution to 100 ng/µL, DNA samples were divided into five pools per group by mixing equally genomic DNA. Pooled DNA samples were randomly sheared into 350 bp fragments, and then end repaired, A-tailed, ligated to paired-end adaptors for PCR amplification. The constructed libraries were sequenced at ∼6.10× coverage on the HiSeq X Ten platform (Illumina, San Diego, USA) by Annoroad Gene Technology Co., Ltd. (Beijing, China).

### Quality control and reads alignment

After removing reads with polluted adaptor, low-quality or over 5% N bases, clean data were carried out on statistics analyses about its quantity and quality. The Burrows-Wheeler aligner was used to map clean reads to the reference genome of *Ovis aries* (Oar v.4.0) (Li and Durbin 2009). Merged alignment files were sorted using SAMtools (Li et al. 2009). Duplicate reads were removed using Picard tools (http://broadinstitute.github.io/picard/) and multiply aligned reads were filtered out.

### Variations calling and annotation

The Genome Analysis Toolkit was used for SNPs and indels calling via local re-assembly of haplotypes for population (McKenna et al. 2010). The SNPs and indels were filtered with following parameters: (SNP: QD < 2.0, ReadPosRankSum < -8.0, FS > 60.0, QUAL < 30.0, DP < 4.0; INDEL: QD < 2.0, ReadPosRankSum < -20.0, FS > 200.0, QUAL < 30.0, DP < 4.0). Then annotation were performed using ANNOVAR for all the qualified variants (Wang et al. 2010). All the called SNPs were filtered to remove loci with low maf (< 0.05) and high missing rate (> 0.10).

### Population genetics analyses

Principal component analysis (PCA) was conducted by EIGENSOFT (Price et al. 2006). Population structure analysis was implemented with Admixture (Alexander et al. 2009). Based on neighbor-joining (NJ) method, PHYLIP was used to construct phylogenetic tree (Plotree D 1989), and the result was displayed by Newick Utilities (Junier and Zdobnov 2010). Linkage disequilibrium analysis were carried out with PopLDdecay (https://github.com/BGI-shenzhen/PopLDdecay).

### Analysis of selective sweep regions

Polymorphism parameters for each breed were calculated by VariScan (Vilella et al. 2005). VCFtools and House-perl scripts were taken to implement fixation index (*F*_ST_) and homozygosity (*H*_P_) respectively, and Z-transformed into *Z*(*F*_ST_) and *Z*(*H*_P_) (Danecek et al. 2011; Axelsson et al. 2013). Both the selective sweep analysis and polymorphism parameters analysis were conducted using a sliding-window method with 100 kb windows sliding at 50 kb steps. Regions with top 5% values of highest *Z*(*F*_ST_) or lowest *Z*(*H*_P_) were recognized as candidate regions.

### Candidate gene analysis

The identified selective regions were annotated to the reference genome (Oar v.4.0), and genes located in selective regions were identified as candidate genes. Gene ontology (GO) analysis was performed in DAVID (https://david.ncifcrf.gov) (Jiao et al. 2012), based on the GO database (http://geneontology.org/) (Harris et al. 2004).

### Data availability

The whole genome sequences of 5 Tibetan sheep breeds are deposited in the NCBI SRA under the accession number SRP266124. The NCBI BioProject ID is PRJNA636698 and BioSample IDs are SAMN15082206 to SAMN15082230. All the supplemental materials have been uploaded in GSA Figshare. Table S1 shows DNA sequencing information of five Tibetan sheep breeds in Qinghai. Table S2 shows SNP and indel calling among five Tibetan sheep breeds in Qinghai. Table S3 shows SNP summary statistics of five Tibetan sheep breeds in Qinghai. Table S4 shows rankings of putative selected regions between different breeds. Table S5 shows lists of putative selected genes between different breeds. Table S6 shows GO analysis of putative selected genes compared with LD breed. Table S7 shows GO analysis of putative selected genes compared with HN breed. Figure S1 shows annotation information of SNPs and indels of Tibetan sheep in Qinghai. Figure S2 shows decay curves of genome-wide linkage disequilibrium and demographic trajectory in Tibetan sheep breeds in Qinghai.

## RESULTS

### Sequencing, alignment, and identification of SNPs

By sequencing 25 DNA pools from five breeds of Tibetan sheep (Figure 1), 2.70 billion reads or 404.56 Gb of genome data were generated and 2.67 billion reads or 401.10 Gb of genome data were yielded via stringent quality filtering for the following analyses (Table S1).

Clean reads were aligned to the reference genome of *Ovis aries* (Oar v4.0) with average coverage rate of 95.00% and average mapping rate of 98.41% (Table S1). A total of 31.36 million SNPs located in chromosomes were identified, and 20.89 million, 20.67 million, 21.19 million, 21.62 million and 21.21 million SNPs were obtained for LD, HN, ZK, TJ and QM respectively (Table S2). Among them, 11.78 million SNPs were shared by five breeds, as well as 1.32 million, 1.08 million, 1.20 million, 1.32 million and 1.17 million SNPs were unique to LD, HN, ZK, TJ and QM respectively (Figure 2A and Table S2). Most SNPs were located in intergenic and intronic regions with T/C replacement, belonged to nonsynonymous/synonymous SNVs (Figure S1A-S1C), Detailed SNP information of each breed was listed in Table S3.

**Figure 2.**
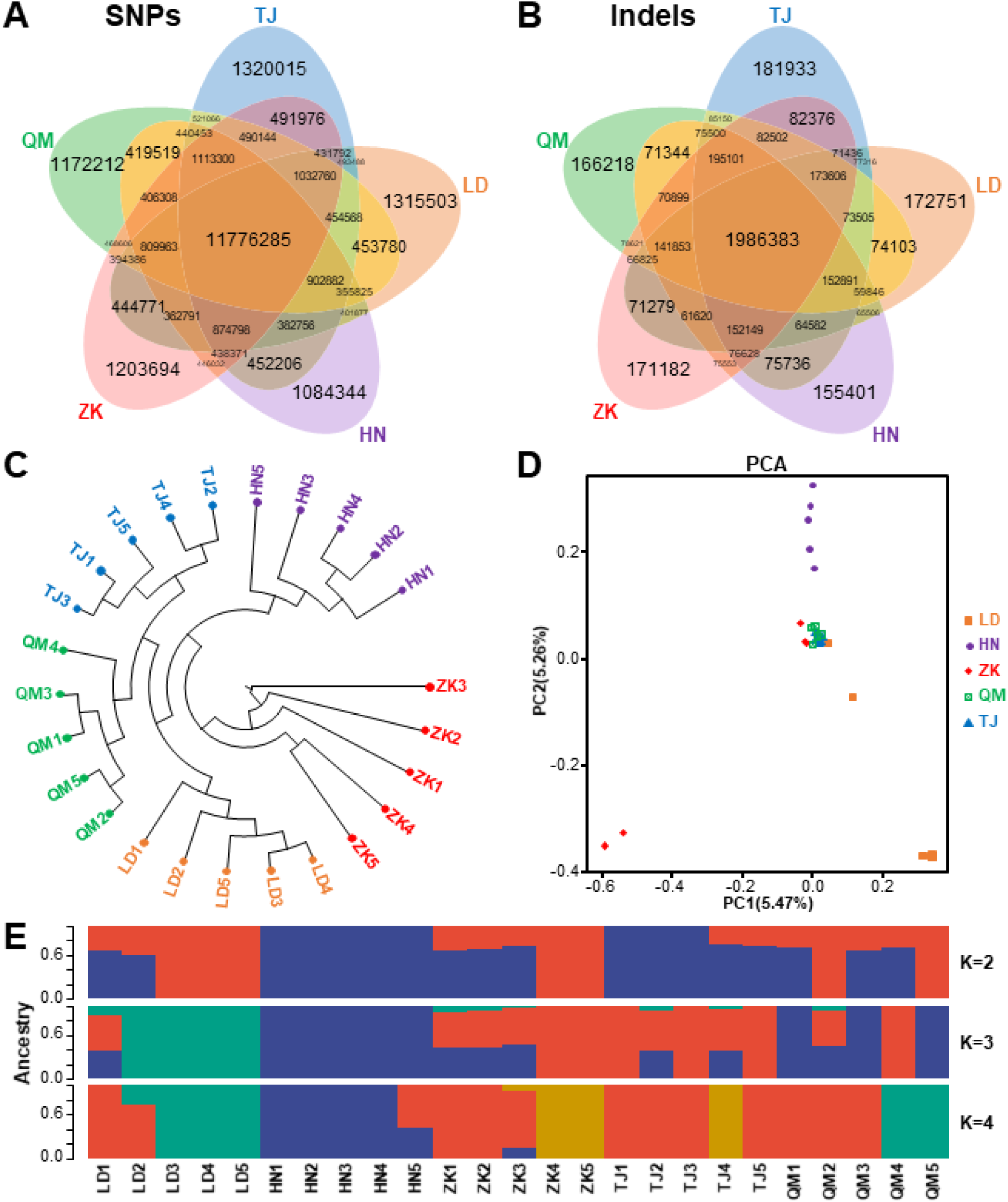
Population genetics analyses of five Tibetan sheep breeds in Qinghai. (**A-B**) Venn diagram showing the shared SNPs (**A**) and indels (**B**) by five Tibetan sheep breeds. (**C-E**) NJ phylogenetic tree (**C**), PCA plots (**D**) and population genetic structure (**E**) of five Tibetan sheep breeds.

In addition, a total of 5.11 million indels were identified with 1.99 million indels shared by five breeds and 0.17 million, 0.16 million, 0.17 million, 0.18 million and 0.17 million indels unique to LD, HN, ZK, TJ and QM respectively (Figure 2B and Table S2). Most indels were located in intergenic and intronic regions with 1 bp length, belonged to frameshift deletion/insertion (Figure S1D-S1F).

### Population genetics structure

To clarify the relationships among Tibetan sheep breeds in Qinghai, a phylogeny tree was constructed using the NJ method. The clustering turned up evidence of separations occurring between breeds, with each breed divided into own clades (Figure 2C). For another, the PCA plotting showed the first two components, explaining 5.47% and 5.26% of the total variation respectively, confirming that QM and TJ were genetically clustered tightly at an intermediate position, while parts of LD and ZK were separated from them respectively. However, the second component clearly differentiated HN from other groups (Figure 2D).

To confirm the degree of divergence, population structure analysis was also implemented to estimate the proportion of common ancestry among breeds at different K values. At K = 2, HN showed a strong genetic differentiation that persisted at higher K values from other groups. At K = 3, LD tended to be separated from the main population in another direction, while other breeds were distributed across the two remaining clusters. When K = 4, a high degree of genetic heterogeneity was observed among breeds, which revealed mixed ancestry. QM possessed a certain genetic similarity with LD, while TJ was much closer to ZK (Figure 2E). In addition, we observed similar decay curves of genome-wide linkage disequilibrium and demographic trajectory in five breeds (Figure S2), suggesting that their habitats were affected by concordant climatic and geologic succession.

### Selective sweep regions related to hypoxic adaptability in Tibetan sheep

Although all the five Tibetan sheep breeds are distributed in Qinghai plateau, their natural habitats are located at different elevations. LD lives below 3000 m, HN, ZK and TJ lives at 3000 to 4000 m, while QM lives over 4000 m. Therefore, we detected selective regions relative to hypoxic adaptability in HN, ZK, TJ and QM breeds using LD as control firstly. There were 167, 193, 163 and 140 selective regions identified for HN (*Z*(*F*_ST_) > 1.83, *Z*(*H*_P_) < -1.90), ZK (*Z*(*F*_ST_) > 1.83, *Z*(*H*_P_) < -1.90), TJ (*Z*(*F*_ST_) > 1.81, *Z*(*H*_P_) < -1.90) and QM (*Z*(*F*_ST_) > 1.80, *Z*(*H*_P_) < -1.90), corresponding to 169, 197, 191 and 158 candidate genes embedded in these selective regions respectively (Figure 3A-3D, Table S4 and S5). Venn diagram showed that 27 identical selective regions (31 genes, including *ABCB7, ADAR, ATP13A3, ATP1B2, CD68, CD74, CYP2E1, DDX60, DIP2C, DNAH2, EIF4A1, FXR2, GP5, KCNN3, MCHR1, MPDU1, MRTFA, NDST1, OR13A1, RPS14, SAT2, SENP3, SH3GL2, SHBG, SOX15, SPRED2, TCOF1, TNFSF12, TNFSF13, TP53, ZMYND11*) were found in 3 breeds, while 4 identical selective regions (7 genes, including *ANXA10, BMP15, CPLANE1, GHR, NIPBL, Olr226, TMEM45A*) were shared by all 4 breeds (Figure 3E and 3F).

**Figure 3.**
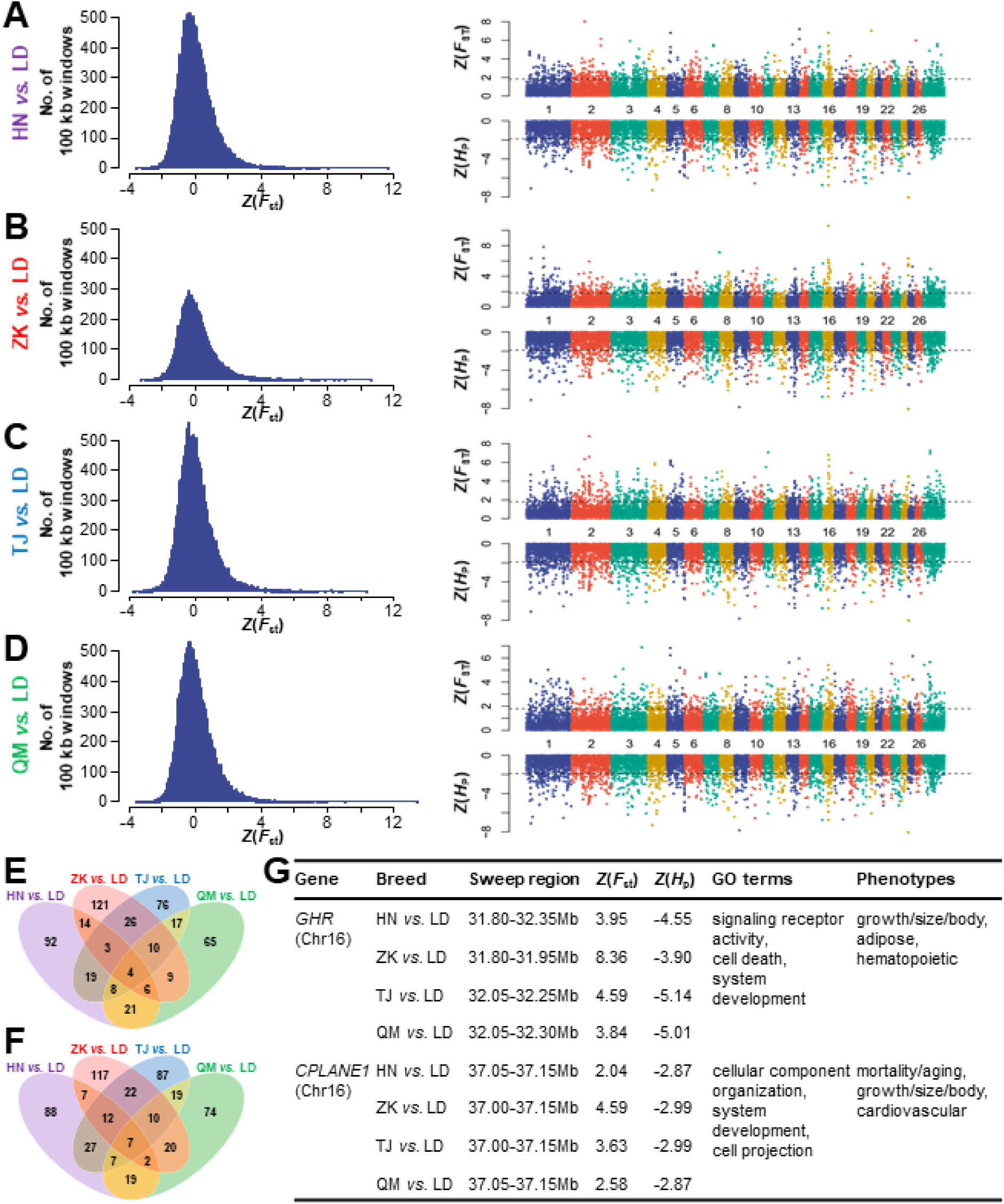
Candidate regions associated with hypoxic adaptability compared with LD breed. (**A-D**) Distribution of average *Z*(*F*_ST_) values for 100 kb windows and plots of *Z*(*F*_ST_) and *Z*(*H*_P_) values along the whole genome in HN *vs.* LD (**A**), ZK *vs.* LD (**B**), TJ *vs.* LD (**C**) and QM *vs.* LD (**D**). A dotted line indicates the cut-off used for extracting outliers. (**E-F**) Venn diagrams of common selected regions (**E**) and corresponding genes (**F**) among different compares. (**G**) Information of representative selective genes in HN, ZK, TJ and QM breeds related to hypoxic adaptability.

To further understand biological functions of selected signals, we performed GO analysis for candidate genes in each breed. We screened selective regions of different breeds relative to LD, 25, 19, 24 and 19 GO terms were significantly enriched (*P* < 0.05) for HN, ZK, TJ and QM respectively. And crucially, genes selected by at least three breeds were enriched in pathways, including membrane, positive regulation of peptidyl-tyrosine phosphorylation, embryonic viscerocranium morphogenesis, positive regulation of histone deacetylation and so on (Table S6). It means that physical development and physiological function of different Tibetan sheep were affected by more severe hypoxia environment, involving in some common important genes such as *CPLANE1* and *GHR* (Figure 3G).

### Selective sweep regions related to coat color in Tibetan sheep

Considering that the physical characteristics of Oula sheep are distinctly different from other breeds, we also detected selective regions in ZK, TJ and QM using HN as control. There were 199, 198 and 146 selective regions identified for ZK (*Z*(*F*_ST_) > 1.81, *Z*(*H*_P_) < -1.89), TJ (*Z*(*F*_ST_) > 1.82, *Z*(*H*_P_) < -1.88) and QM (*Z*(*F*_ST_) > 1.82, *Z*(*H*_P_) < -1.88), corresponding to 206, 223 and 157 candidate genes respectively (Figure 4A-4C, Table S4 and S5). These selective regions were intersected and 15 selective regions (24 genes, including *AP2B1, BICD2, BSND, ERMP1, FGD3, GAS2L2, IPPK, LCORL, MAP6D1, NCAPG, PARL, PFN2, POGZ, PSMB4, R3HDM1, RASL10B, RIC1, SELENBP1, SUPT3H, TAFA1, TMEM61, USP24, YEATS2, ZC4H2*) were shared by them (Figure 4D and 4E).

**Figure 4.**
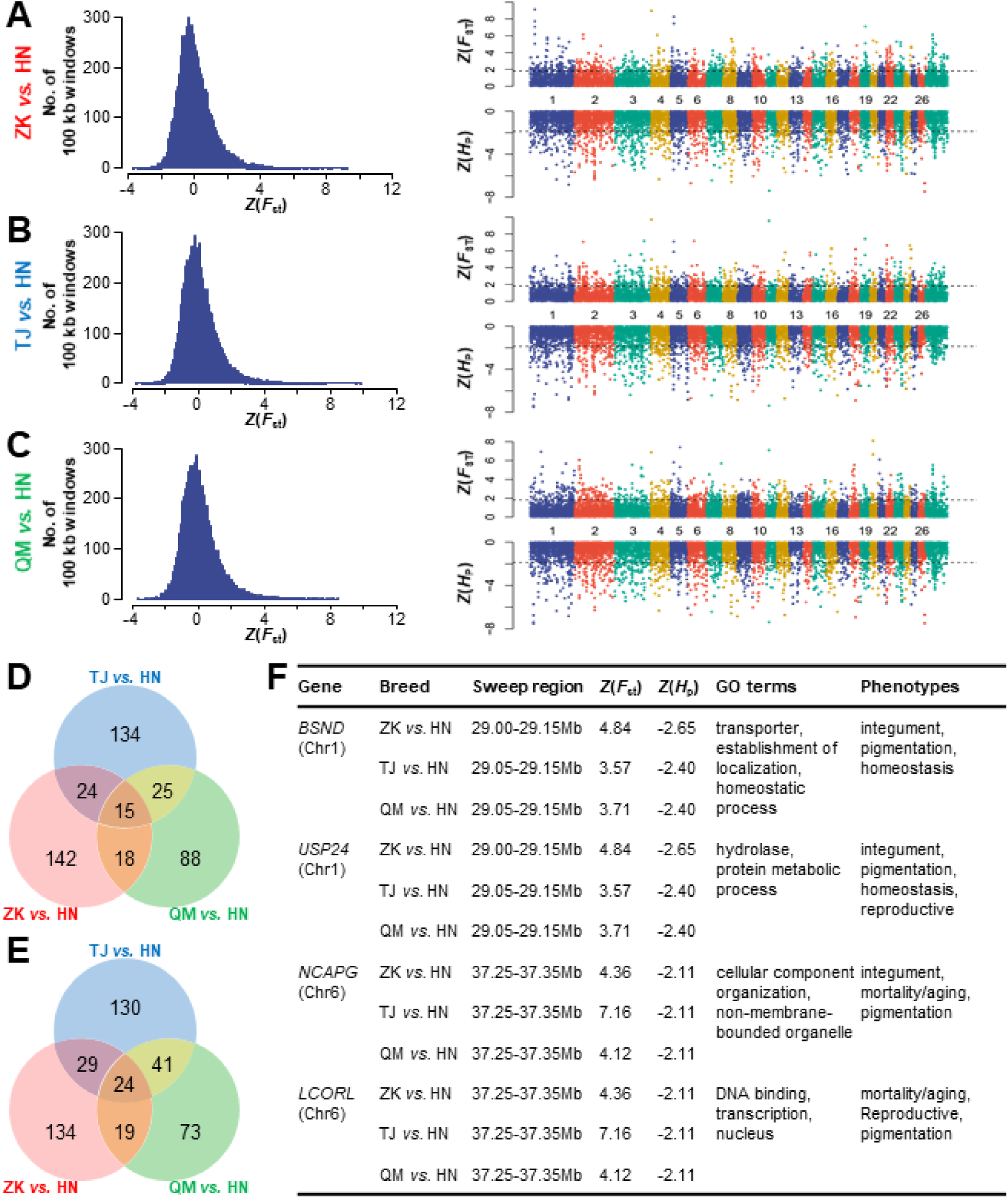
Candidate regions associated with physical characteristic compared with HN breed. (**A-C**) Distribution of average *Z*(*F*_ST_) values for 100 kb windows and plots of *Z*(*F*_ST_) and *Z*(*H*_P_) values along the whole genome in ZK *vs.* HN (**A**), TJ *vs.* HN (**B**) and QM *vs.* HN (**C**). A dotted line indicates the cut-off used for extracting outliers. (**D-E**) Venn diagrams of common selected regions (**D**) and corresponding genes (**E**) among different compares. (**F**) Information of representative selective genes in ZK, TJ and QM breeds related to physical characteristic.

GO analysis for candidate genes showed 12, 21 and 16 GO terms were significantly enriched (*P* < 0.05) for ZK, TJ and QM relative to HN respectively. Furthermore, common selected genes in three compares were closely related to cytoskeleton and negative regulation of microtubule depolymerization (Table S7). Interestingly, numerous selective genes, including *BSND, USP24, NCAPG* and *LCORL*, are found to be related to integument, pigmentation, homeostasis and mortality (Figure 4F). This might explain why the trait of Oula sheep is vastly different from other Tibetan sheep breeds.

## DISCUSSION

Tibetan sheep is one of the three original sheep breeds in China, which formed and evolved with northern Chinese migration (Hu et al. 2019). Through long-term natural selection, Tibetan sheep are well suited to the harsh climate variation and poor pasture condition of plateau region, associated with morphological, physiological and genetic changes (Niu et al. 2016; Jing et al. 2019). Similar phenomena also have been found in other indigenous herbivores, such as yak (Qiu et al. 2012), Tibetan antelope (Rong et al. 2012) and gazelles (Zhang and Jiang 2006). Although Tibetan sheep genome is also deciphered to illuminate its evolutionary history and biological peculiarity (Hu et al. 2019), mechanism about its adaptability dealing with different environments remains puzzling. Here we compared genomic changes of five Tibetan sheep breeds, and confirmed that Tibetan sheep under different circumstances have formed own branches and generated distinctive traits.

Hypoxia is the most striking feature of environmental changes in Qinghai-Tibet plateau (Ji et al. 2016). Almost all of plateau mammals have evolved the high-altitude adaptability to adjust physiological function to accommodate the hypoxic environment. Despite of big differences in genotype and phenotype between animals living in plain and plateau, Tibetan sheep improve the progressive tolerance to hypoxia with a rising of living altitude (Wei et al. 2016). Through comparative genomic analysis, we found a series of SNPs and corresponding candidate genes concerned with hypoxic adaptability of Tibetan sheep at different regions. Among them, *GHR* encodes a transmembrane receptor for growth hormone, and has been closely related to the adaptability and growth traits of Tibetan sheep (Ma 2007; Han et al. 2016). *BMP15* is also an important gene encoding a secreted ligand of the transforming growth factor-β and affecting the expression of downstream transcription factors. It’s been widely reported that mutations happened in *BMP15* gene are associated with prolificacy and reproduction traits of sheep (Abdoli et al. 2018; Dolebo et al. 2019). *CPLANE1* is a new gene we found that could be associated with physiological function of Tibetan sheep in higher altitude. It encodes a transmembrane protein responsible for mitosis and neurogenesis, which defects could be a cause of Joubert syndrome(Hong et al. 2019; Srour et al. 2012). Moreover, some different genes were positively selected in connection with hypoxic adaptation in different comparison, such as *TP53* in HN, ZK and TJ (Muller et al. 2019), *MCHR1* in ZK, TJ and QM (Diniz and Bittencourt 2019).

Different from other Tibetan sheep breeds, Oula sheep tends to have a larger physique with sparse and dried brown wools, and its early development and meat performance are better than others (Liu et al. 2015). Some reports suggested that Oula sheep is originated from hybridization of wild argali and Tibetan sheep (Xian et al. 2017). We identified several genes responsible for this phenotypic modulation, especially coat color, including *BSND, USP24, NCAPG* and *LCORL*. Except for integument, BSND mutation is also correlated with failure to thrive and decreased body size (de Pablos et al. 2014; Nomura et al. 2011). *NCAPG* is required for the condensation and stabilization of chromosomes during mitosis and meiosis (Murphy and Sarge 2008),which regulates proliferation and apoptosis in carcinoma cells (Liu et al. 2018). Polymorphisms in *LCORL*, an important transcription factor, are related with skeletal frame size and adult height (Horikoshi et al. 2013).

## CONCLUSION

In this study, we identified a novel series of genes and function mutations subjected to positive selection in different breeds of Tibetan sheep. Hypoxic response of Tibetan sheep living in higher altitude area was related with genes including *BMP15, GHR* and *CPLANE1*. Furthermore, some specific selective genes, such as *BSND, USP24, NCAPG* and *LCORL*, may explain why Oula sheep is distinctly different from other breeds in coat color, size appearance and growth property. These results provide new insights into the molecular mechanism of Tibetan sheep domestication and evolution as well as the formation of the unique characteristics of different Tibetan sheep breeds.

## ACKNOWLEDGEMENTS

This study is funded by the Strategic Priority Research Program of Chinese Academy of Sciences (XDA2005010406) and the Natural Science Foundation of Qinghai Province (2017-ZJ-915Q and 2017-NK-114). GJ is supported by “CAS Light of West” programs and HF is supported by Qinghai “1000 Talents” programs. We thank Ledu, Henan, Zeku, Qumarlêb ang Tianjun Animal Husbandry and Veterinary Station for the help with animal sampling.

---

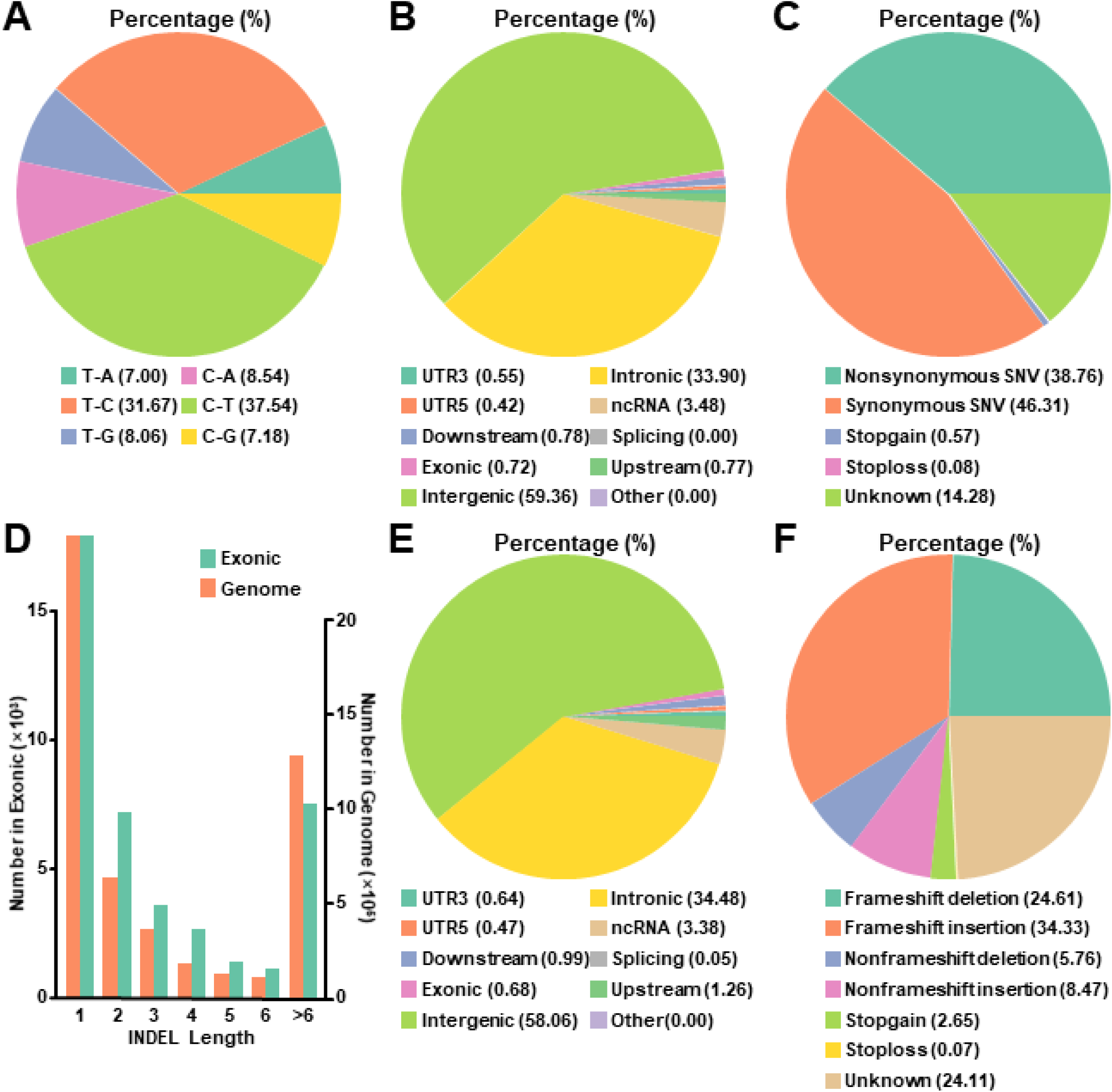

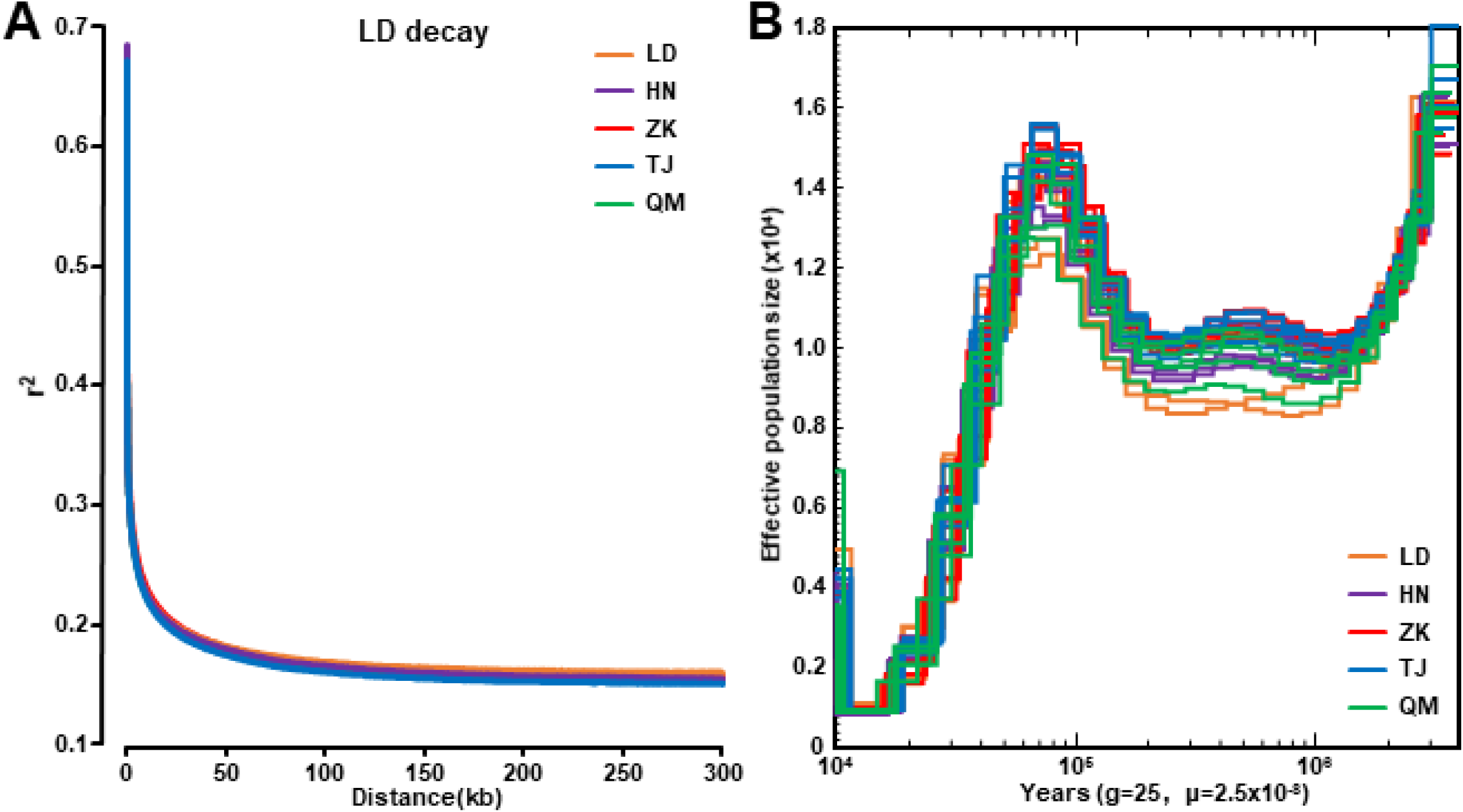

